# Application of new informatics tools for identifying allosteric lead ligands of the c-Src kinase

**DOI:** 10.1101/038323

**Authors:** Lili X. Peng, Morgan Lawrenz, Diwakar Shukla, Grace W. Tang, Vijay S. Pande, Russ B. Altman

## Abstract

Recent molecular dynamics (MD) simulations of the catalytic domain of the c-Src kinase revealed intermediate conformations with a potentially druggable allosteric pocket adjacent to the C-helix, bound by 8-anilino-1-naphthalene sulfonate. Towards confirming the existence of this pocket, we have developed a novel lead enrichment protocol using new target and lead enrichment software to identify sixteen allosteric lead ligands of the c-Src kinase. First, Markov State Models analysis was used to identify the most statistically significant c-Src target conformations from all MD-simulated conformations. The most statistically relevant candidate MSM targets were then prioritized by assessing how well each reproduced binding poses of ligands specific to the ATP-competitive and allosteric pockets. The top-performing MSM targets, identified by receiver-operating curve analysis, were then used to screen the ZINC library of 13 million ‘clean, drug-like’’ ligands, all of which prioritized based on their empirical scoring function, binding pose consistency across MSM targets, and strong hydrogen bonding and hydrophobic interactions with Src residues. The FragFEATURE knowledgebase of fragment-protein pocket interactions was then used to identify fragments specific to the ATP-competitive and allosteric pockets. This information was used to identify seven Type II and nine Type III lead ligands with binding poses supported by fragment predictions. Of these, Type II lead ligands, ZINC13037947 and ZINC09672647, and Type III lead ligands, ZINC12530852 and ZINC30012975, exhibited the most favorable fragment profiles and are recommended for further experimental testing for the existence of the allosteric pocket in Src.

## INTRODUCTION

Drug discovery is a complex and difficult processes in the pharmaceutical industry as millions of dollars and man-hours are devoted to the discovery of new therapeutic agents. Traditional methods of drug discovery rely on trial-and-error testing of chemical substances on *in vitro* and *in vivo* biological models, and matching apparent effects to treatments.^1^ A commonly used computational method to identify promising compounds to bind to a target molecule of known structure is virtual screening. There are two main approaches of virtual screening: structure-based approaches, which involve docking of candidate ligands into a protein target followed by applying a scoring function to estimate the affinity of the ligand for the protein, and ligand-based approaches, which rely on knowledge of known binders to create a pharmacophore model for the target of interest.^2^ Virtual screening is a productive and cost-effective way to search for novel lead compounds, especially given the increasing availability of high-performance computing platforms. However, the accuracy of algorithms used in docking software still has room for improvement. In addition, the design of lead compounds also heavily relies on the amount and quality of structural information on the target protein.^3^ A protein exists in a population of conformational states in dynamic equilibrium with one another, adopting conformations not always captured by X-ray crystallography and NMR spectroscopy.^4^,^5^Large-scale supercomputing platforms such as Anton^6^, Google Exacycle^7,8^, and Folding@Home^9^ have been used to run molecular dynamics (MD) simulations on proteins with important disease implications, such as the epidermal growth factor receptor kinase^10^ and G-protein coupled receptor^7^, to reach timescales on the order of hundreds of microseconds. These simulation results reveal conformational states not yet witnessed experimentally and could be potentially exploited for drug design.^11-14^

The proto-oncogene tyrosine-protein kinase Src plays important roles in cell proliferation, migration, and survival.^13-16^ Recent ∽50 µs massively-distributed parallel MD simulations on the catalytic domain of c-Src reveal a hydrophobic pocket, adjacent to the ATP binding site and encapsulated by the αC-helix, β4, and β5 strands, that is structurally similar (21% identity) to hydrophobic allosteric pocket in cyclin-dependent kinase 2 (Cdk2), bound with 8-anilino-1-naphthalene sulfonate (ANS).^17^ The binding of ANS to Cdk2 is accompanied by substantial structural changes, inducing αC-helix conformation incompatible with association of Cdk2 with downstream substrate cyclin A.^16^ Superimposition of representative simulation snapshots of ANS-bound and *apo* ANS conformations of intermediate Src shows that ANS binding displaces the αC-helix outwards in a manner similar to how binding of ANS displaces the αC-helix in Cdk2 (see Figure 1). The binding of ANS remains energetically stable throughout the ∽50 µs aggregate MD simulations, suggesting that the ligand can bind to the pocket in a manner similar to ANS binding Cdk2. Figure 1 shows that the binding pose of ANS blocks the formation of the salt bridge between the catalytic Lys295 and Glu310. This salt bridge is necessary for Src activation.^16^,^18^Likely additional contributions to ANS’ binding pose include hydrogen bonding with the side-chain amine group of Lys295 and backbone heavy atoms of Phe405 and Gly406, as well as ample hydrophobic interactions between naphthalene and phenyl rings with neighboring hydrophobic residues.^19^ Altogether these findings strongly suggest the presence of an allosteric pocket in MSM states of intermediate Src conformations that could be exploited for discovery of allosteric Src inhibitors.

**Figure 1.**
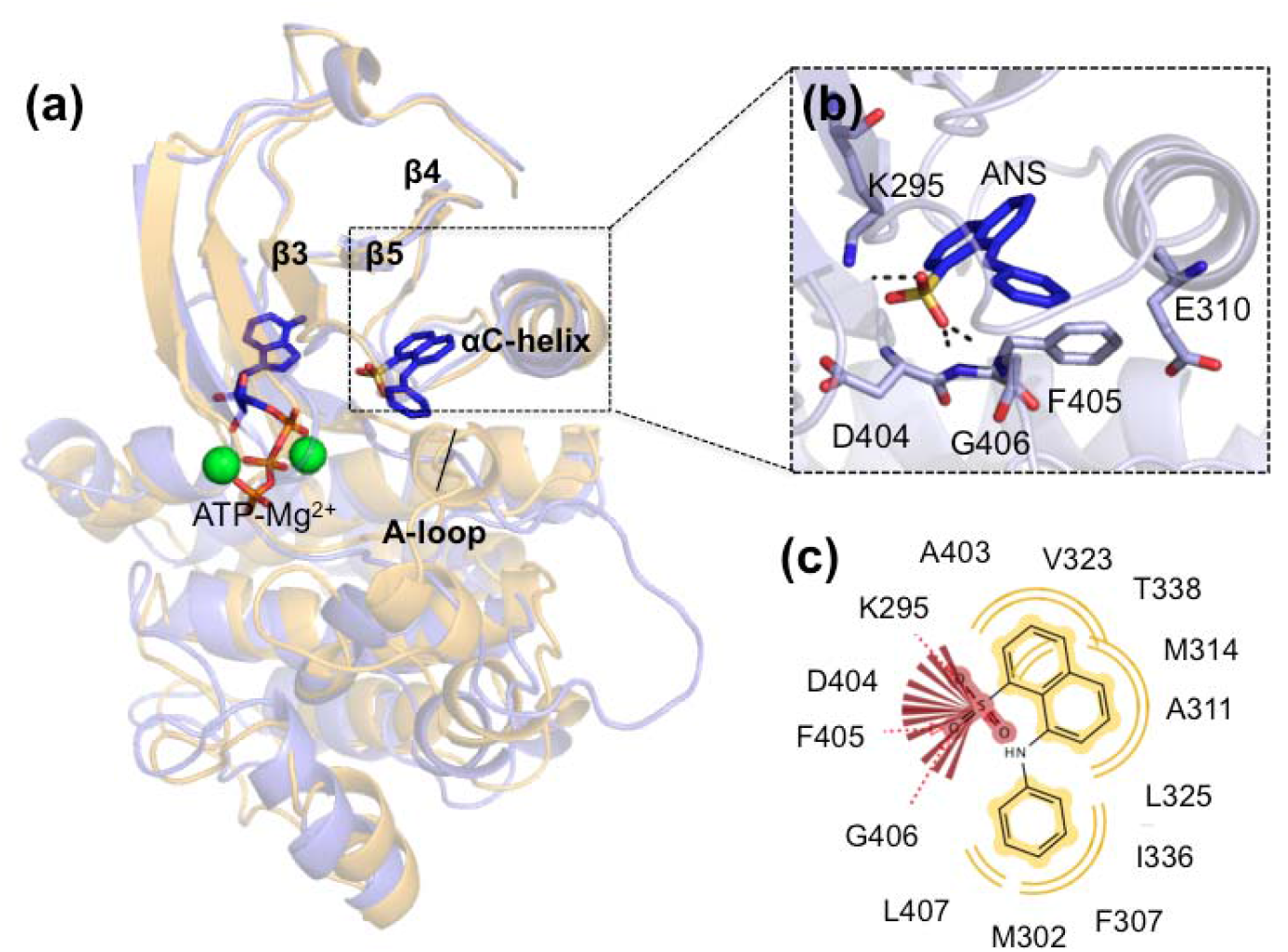
a) Local superimposition of the *apo* ANS (pink) and ANS-bound (blue) intermediate Src conformations reveals that ANS binding displaces the αC-helix outwards relative to αC-helix’s orientation in the *apo* ANS MSM state. The allosteric pocket is encapsulated by β4, β5, and the αC-helix. b) Binding of ANS can be stabilized by hydrogen bonding of the sulfonate group to the Lys295 sidechain amide and backbone amides of Asp404, Phe405, and Gly406, thereby preventing formation of the salt bridge between Lys295 and Glu310. c) Pharmacophore of ANS shows hydrogen bonding with Lys295 and DFG residues (red) and hydrophobic interactions between the naphthalene ring and Val323, Thr338, Met314, Ala311, and Leu325 and between the phenyl ring and Leu407, Met302, Phe307, Ile336, and Leu325.

Allosteric targeting of kinases via inhibitor binding has been gaining traction in drug discovery because they overcome target selectivity by exploiting non-ATP-competitive binding sites and regulatory mechanisms unique to a particular kinase.^20^-^23^ The objective of this study is to recommend lead ligands specific to this allosteric pocket in Src for experimental testing. Towards this end, we developed a generalizable workflow using new predictive analytics software for target and lead enrichment, MSMBuilder^24^ and FragFEATURE^25^, respectively, and using qualitative metrics for various enrichment steps. This workflow was applied to determine ligands that bind the allosteric and ATP-competitive pockets (Type II) and those that bind strictly the allosteric pocket (Type III). We foresee that these predicted ligands could be used as small molecules to experimentally procure a co-crystal structure of an intermediate Src conformation, to prove the existence of the allosteric pocket.

It is noteworthy that, in generating candidate target conformations of Src, we had considered using less computationally intensive approach like homology modeling, which would involve stringing c-Src’s sequence onto the crystal structure of ANS-bound Cdk2 as a template. However, clustal sequence alignment of human Src and Cdk2 returns 21% identity and 35% similarity, based on the kinase domain, values too low to have confidence in homology modeling.^26^ Our approach of running long-timescale MD simulations utilizes a previously generated MD dataset that is derived from the experimental crystal structures of the inactive and active conformations of the Src kinase, followed by MSM analysis, which yields mechanistic insight into the protein-ligand system within a set of physics-derived parameters of an MD force field.^27^ Altogether these features offer a very rigorous approach in identifying target conformations of Src for virtual screening, an advantage over less computationally demanding approaches like homology modeling. Furthermore, the application of Markov State Models to detect intermediate conformations and cryptic allosteric binding pockets has been gaining increasing user adoption in the field of protein conformational change. Buch et al. used MSMs ti predict the binding pose and binding pathways of trypsin.^28^ Plattner et al. have also MSMs to understand conformational change and ligand-binding kinetics for trypsin and its inhibitor benzamidine.^29^ Finally, Greg Bowman and colleagues have also used MSMs to detect allosteric sites in β-lactamase, interleukin-2, and RNase H.^30^,^31^

This workflow commences with performing MSM analysis on all conformational states accessed during the MD simulations, followed by selecting candidates based on quantitative metrics, such as statistical significance and the binding pose reproducibility of ATP-Mg^2+^ and ANS, and qualitative metrics, such as the size and morphology of the software-defined docking region. The best-performing target conformations were then used to screen the ZINC library of 13 million “clean, drug-like” ligands.^32^ After selecting the lead ligands with the highest docking scores, they were further prioritized by whether they were bound to the appropriate binding region (Type II: ATP-competitive and allosteric, Type III: allosteric) and consistency in binding pose across multiple target conformations. Furthermore, Figure 1 shows that ANS binding Src could be stabilized by hydrogen bonding to the side chain amide of Lys295 and backbone heavy atoms of the DFG motif. The naphthalene ring of ANS could interact with hydrophobic residues Ala311, Met314, Val323, Leu325, and Thr338; the phenyl ring of ANS could interact hydrophobically with Met302, Phe307, Leu325, Ile336, and Leu407.^17^ These findings motivated us to prioritize lead ligand candidates exhibiting hydrogen bonding and hydrophobic interactions similar to those of ANS-bound Src.

The workflow culminates with identification of lead ligands whose substructures are empirically supported to bind to pockets (of other proteins) structurally similar to the docking region of Src target conformations. This information on protein structural environments annotated with small molecule fragments is in the FragFEATURE knowledge base, curated from 34,000 Protein Data Bank structures using *k* nearest neighbors, a supervised machine learning algorithm.^25^ FragFEATURE compares structural environments from a target protein to the knowledgebase with similar structural environments and identifies statistically preferred fragments. FragFEATURE was developed to identify fragments that could lead to drug-like ligands to study further using computational and/or experimental tools in lead discovery, using a data-driven approach given the availability of the large number of structures of protein-small molecule complexes. Currently, conventional methods of *in silico* fragment-based binding predictors/evaluators rely on physics-based scoring functions, which can be difficult to calibrate. FragFEATURE circumvents these limitations by making predictions based on the statistics and direct observations of fragment contacts with amino acid side chains and their environments. FragFEATURE is entirely data-driven and predicts the likely binding of a fragment because the environment in question looks very similar to other environments where that fragment has been observed to bind previously.^33^

The study describes how new computational tools, Markov State Models and FragFEATURE, can be used to enrich target conformations from long timescale MD simulations and reduce the chemical search space in lead identification. Markov State Models have been previously used in drug discovery to identify the prominent target conformations of GPCRs^7^, but this work more importantly focuses on enrichment of lead ligands, using the FragFEATURE knowledge base of preferred fragment-protein pocket interactions. This study represents the first time FragFEATURE is used in lead enrichment; interpretations of the FragFEATURE results are discussed in the context to identify lead candidates with the strongest fragment profiles, and subsequently recommended for experimental testing. Overall, this methodology has potential to be incorporated in lead enrichment protocols in early stages of drug discovery.

## METHODS

### Enrichment of MSM states of intermediate Src conformational targets

The first step of target enrichment was to optimize the number of candidate target conformations from 97983 MD snapshots from Shukla et al.’s MD simulations of ANS-bound intermediate Src totaling 50 μs.^17^ Screening all snapshots would be computationally intractable, so MSMs were constructed to identify the dominant conformational states of ANS-bound intermediate Src. This was achieved by clustering all snapshots by residues within 5 Å of ATP-Mg^2+^ and ANS at an RMSD cutoff of 2.6 Å, after testing a range of values. This cutoff selection of 2.6 Å returned ten MSM conformational states A-J, which sufficiently captures the diversity of the full dataset (see Figure 2 and Table 1) and are conformationally representative of the 97983 MD snapshots. While we could have used the traditional protein clustering method with an RMSD cutoff to identify conformational states, this method provides structurally distinct conformations with insignificant equilibrium populations. On the other hand, MSMs helps us identify conformational states with highest equilibrium population from a set of simulation trajectories.

**Table 1.**
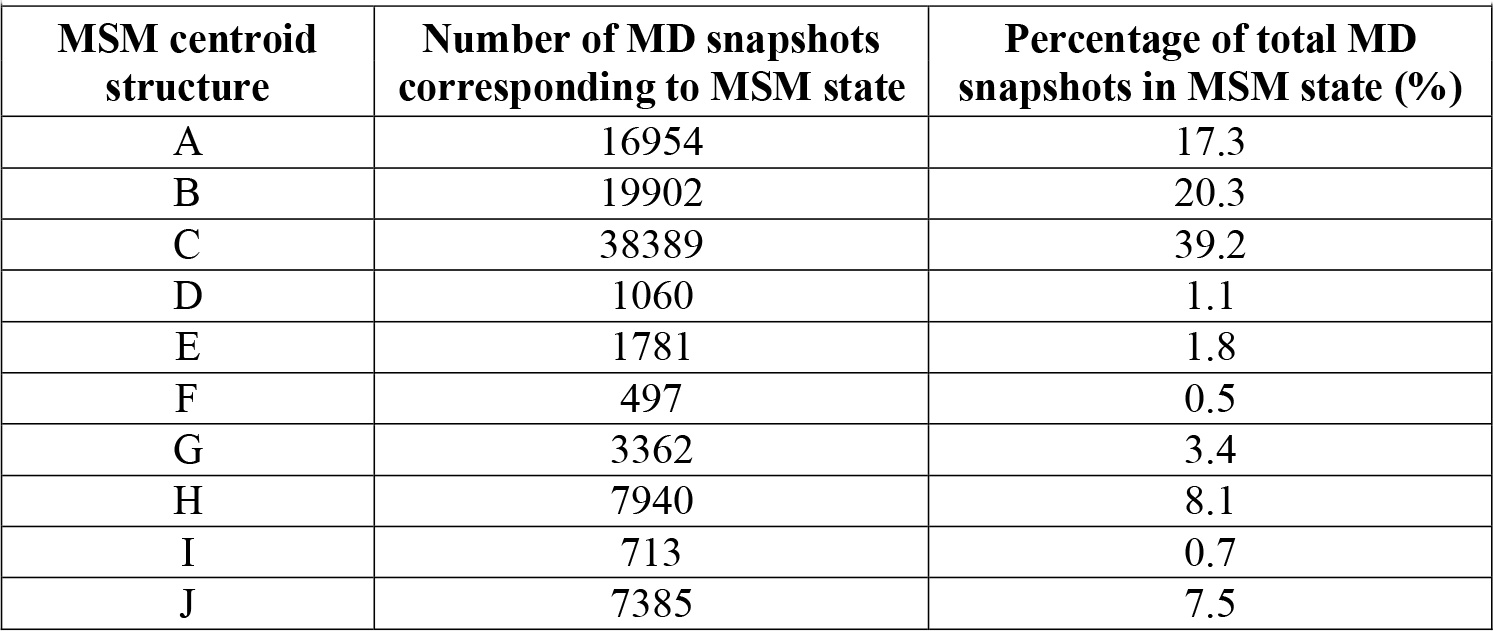
Statistical significance of MSM states of *k =* 10 centroid structures.

**Figure 2.**
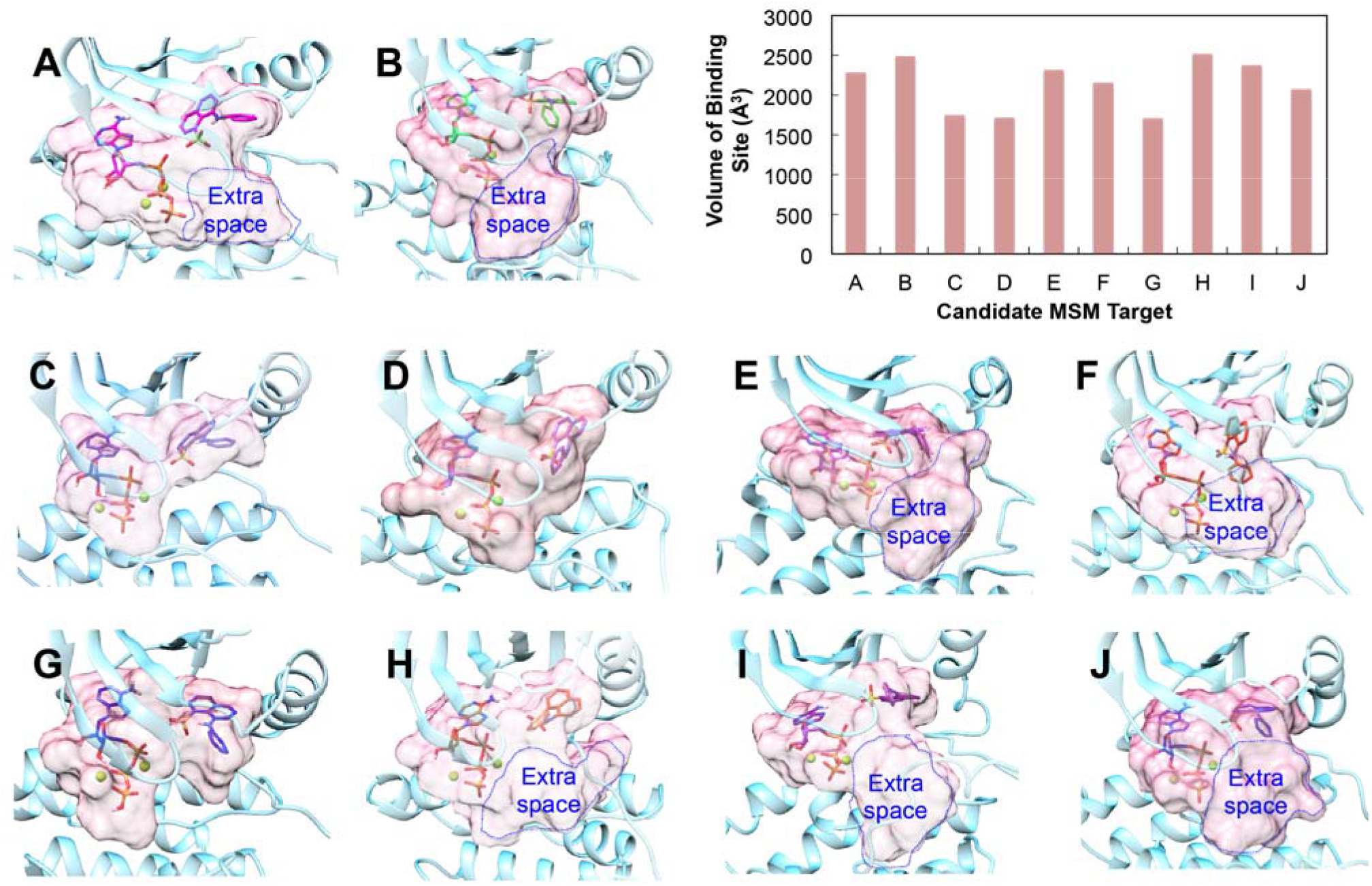
Size and morphology of hybrid ATP-Mg^2+^, ANS-derived binding sites in the ten candidate MSM targets. Upper right-hand inset shows the volume of the binding site for each candidate target: A (2286 Å^3^), B (2489 Å^3^), C (1747 Å^3^), D (1717 Å^3^), E (2314 Å^3^), F (2154 Å^3^), G (1705 Å^3^), H (2513 Å^3^), I (2374 Å^3^), and J (2107 Å^3^).

The next step was to screen for ligands that would dock the ATP-competitive and allosteric regions (Type II) or strictly the allosteric region (Type III), the docking area of each candidate target was derived from the original binding poses of ATP-Mg^2+^ and ANS, respectively, to designate a ‘hybrid’ docking area of each MSM target. Using Surflex-Dock^34^, the threshold, which determines how much of the binding site can be buried in the protein, and bloat, which is used to inflate the docking area and nearby crevices, were set to default values of 0.5 and 0.0, respectively. All ligands were docked invoking the standard scoring function (pscreen) and receptor proton flexibility (pflex).

The docking area in our MSM states of intermediate Src conformations includes the ATP-competitive and allosteric pockets. Prior to commencing target enrichment, we used the receiver-operating characteristic (ROC) approach to evaluate our docking method in its ability to distinguish between the active and inactive (decoy) ligands to a specific site on the target.^35^ While this method is applicable for the ATP-competitive binding site, which has known active and decoy ligands, the allosteric pocket is witnessed only in the presence of ANS binding intermediate Src conformations from MD simulations. Because there are currently no publicly accessible active and decoy ligands specific to the allosteric site, the ROC approach is unsuitable for testing the performance of the allosteric region. However, for benchmarking against the intermediate Src conformations, the ROC approach was applied to test performance of the ATP-competitive pocket: the Directory of Useful Decoys (DUD)^36^ has 126 active ligands for AMPCPP, an ATP mimetic in which the oxygen between P_α_ and P_β_ is mutated to carbon, which is bound to an inactive, DFG-in conformation of Src (PDB id 2SRC^37^). In addition to using ROC for the ATP-competitive region, we sought to define other criteria to prioritize the ten candidate MSM targets. The first criterion we had used is statistical significance of each MSM centroid state, or the number of corresponding MD snapshots with respect to the total number of MD snapshots (97983) from the aggregate 50 microsecond MD simulation. Targets A, B, and C exhibited the highest statistical significance in its representation of the protein conformational space, at 17.3%, 20.3%, and 39.2%, respectively (see Table 1). While statistical significance is important in target enrichment, visual inspection of the docking regions in the ten candidate targets reveal considerable differences in their three-dimensional size (volume) and morphology. Figure 2 shows that docking regions for targets C, D, and G exhibit relatively conservative coverage of the ATP-competitive and allosteric sites. This is indicated by the docking region volumes of 1747 Å^3^, 1717 Å^3^, and 1705 Å^3^, which are below the average volume of 2141 ± 315 Å^3^. On the other hand, the allosteric pocket in other targets, including A and B, are slightly larger in that they occupy space not derived from ATP-Mg^2+^ and ANS. While these differences in the size and morphology of a target’s docking region seem minor, it is likely that they can result in significant differences in docking scores of a ligand’s predicted binding pose. Finally, another criterion used to select target conformations for virtual screening is how well each MSM target reproduces the binding poses of ATP-Mg^2+^ and ANS. How these criteria are used all together to prioritize the ten candidate MSM states is discussed in more detail in the succeeding sections.

*Applying the ROC approach to the ATP-competitive region:* ROC curves for each of the ten candidate MSM targets were calculated, along with area-under-the-curve (AUC) values, using the 126 active ligands for AMPCPP bound to inactive Src (PDB id 2SRC) from DUD^36^ and 990 randomly selected decoy ligands from Bissantz et al.^38^ The average AUC is 0.65 ± 0.03, with targets A and G having the highest and lowest AUC values of 0.72 and 0.61 respectively. These values are not particularly outstanding but consistent with the standard AUC of ∽0.6 for kinases.^35^ However, the highest AUC of 0.72 (from target A) is nearly equivalent to 0.73, the AUC for the inactive, AMPCPP-bound Src conformation (2SRC), as shown in Figure S4, suggesting that the best-performing target, in terms of AUC, is consistent with the performance of the crystal structure of the inactive conformation.”

*Three-dimensional size and morphology of the hybrid ATP-Mg^2+^, ANS-derived binding site:* As aforementioned, Figure 2 shows that the docking regions of targets C, D, and G exhibit sufficient but conservative coverage of the ATP-competitive and allosteric sites, whereas the docking regions in the other MSM targets extend outwards of the allosteric pocket towards the A-loop, tangential to the region occupied by the phosphate of ATP-Mg^2+^ and the region occupied by ANS. The docking regions of targets C, D, and G are, on average smaller (1723 Å^3^), than those of the other seven candidates (2320 Å^3^). This region extending towards the A-loop may seem extraneous, since it is neither derived from the binding poses of ATP-Mg^2+^ and ANS, and could introduce undesirable poses in ligands during virtual screening. However, later we were alerted to Type III allosteric inhibitors of MEK1/2 and p38α: visual inspection of PD318088-bound MEK1/2^39^ (PDB id 1S9J) and GDC-0973-bound p38α^40^ (PDB id 4AN2) reveal allosteric regions that also extend towards the A-loop. The allosteric pockets in MEK1/2 and p38 are structurally similar to that in Cdk2 and MSM targets of ANS-bound intermediate Src, as evidenced by their structural overlay in Figure S1. This evidence strongly suggests that the region extending towards the A-loop in intermediate Src conformations could be exploited for enhancing Src selectivity. Not present in targets C, D, and G, this space is present in docking regions of targets A, B, E, F, H, I, and J, all together representing 56.2% of 97983 trajectory snapshots.

*Binding pose reproducibility of ATP-Mg^2+^ and ANS:* Assessment of how well each MSM target reproduces the binding poses of ATP-Mg^2+^ and ANS was done by redocking each ligand into its original target, and calculating the RMSD between the original and re-docked poses for the three top-ranking poses (see Tables 2 and S1). Good reproducibility was defined by an RMSD not exceeding 3 Å. Based on this cutoff, targets A, E, G, and H reproduce the binding pose of ATP-Mg^2+^ well, and targets A, F, and J exhibit for ANS. Notably, of the ten candidate targets, only target A exhibits good reproducibility for both ATP-Mg^2+^ and ANS (RMSD of 2.35 ± 1.82 Å for ATP-Mg^2+^, RMSD of 2.34 ± 0.31 Å for ANS). In the context of target selection, heavier emphasis was placed on the binding pose reproducibility of ANS than that of ATP-Mg^2+^ because there are no known active ligands to the allosteric pocket of Src, leaving ANS’ binding pose reproducibility the only quantitative metric for assessing performance of the allosteric region in each candidate target.

**Table 2.**
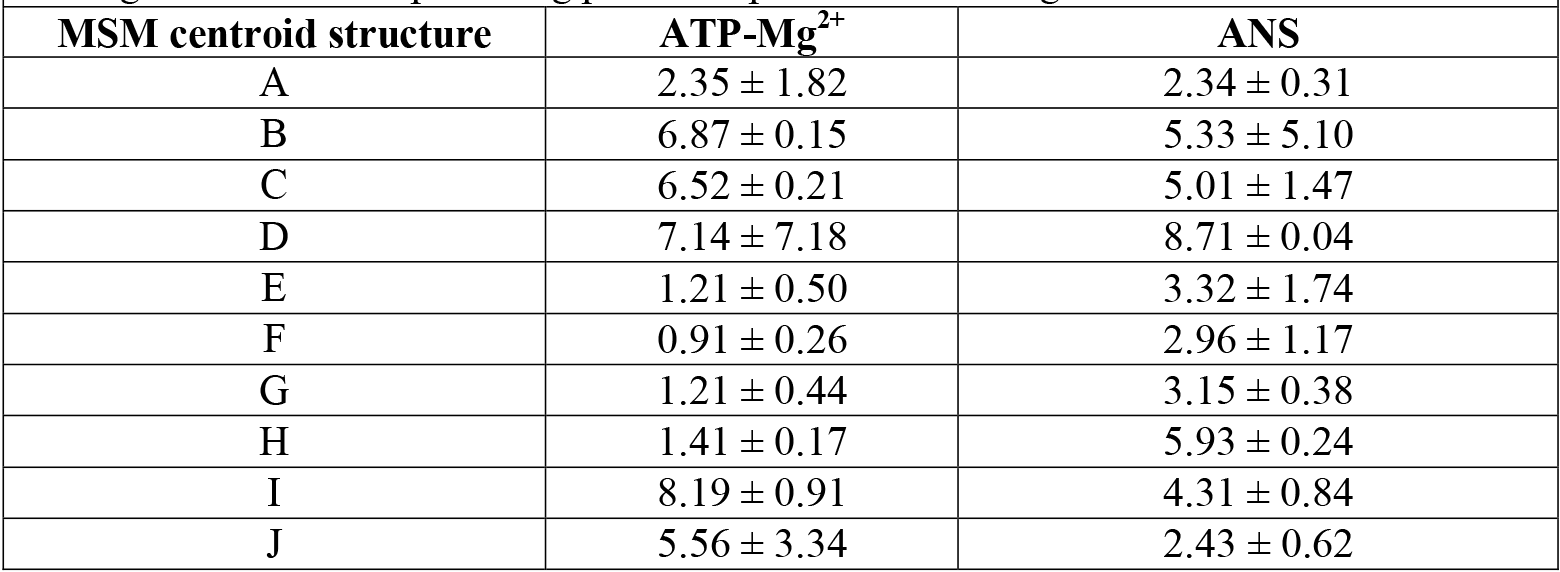
Binding pose reproducibility by average RMSD (Å) of original and predicted poses of ATP-Mg^2+^ and ANS of *k =* 10 centroid structures. Values were calculated from the original and three top-ranking predicted poses for ATP-Mg^2+^ and ANS.

**Table 3.** Type II and III lead ligand candidates with good to excellent binding pose consistency between MSM targets A and B. Shows the ZINC ID number, molecular weight, chemical formula, RMSD between binding poses in MSM targets A and B.

| Type II lead ligand candidates |  |  |  |
| --- | --- | --- | --- |
| ZINC ID | Molecular Weight (g/mol) | Chemical Formula | Pose RMSD between targets (Å) |
| ZINC13037947 | 444.6 | $C_{22}H_{28}N_4O_4S$ | 2.343 |
| ZINC09672647 | 488.6 | $C_{23}H_{28}N_4O_6S$ | 1.955 |
| ZINC15729866 | 495.6 | $C_{25}H_{29}N_5O_4S$ | 2.533 |
| ZINC39795560 | 464.5 | $C_{24}H_{24}N_4O_4S_1$ | 2.203 |
| ZINC12939211 | 418.4 | $C_{18}H_{18}N_4O_6S$ | 1.677 |
| ZINC09125889 | 468.6 | $C_{22}H_{24}N_6O_2S_2$ | 2.247 |
| ZINC09276644 | 429.5 | $C_{22}H_{24}FN_3O_3S$ | 2.584 |
| Type III lead ligand candidates |  |  |  |
| ZINC ID | Molecular Weight (g/mol) | Chemical Formula | Pose RMSD between targets (Å) |
| ZINC55183050 | 320.4 | $C_{21}H_{23}NO_2$ | 2.541 |
| ZINC14369395 | 382.4 | $C_{21}H_{19}FN_2O_4$ | 2.544 |
| ZINC80660181 | 347.4 | $C_{17}H_{26}N_6O_2$ | 1.214 |
| ZINC12543756 | 408.4 | $C_{22}H_{21}FN_4O_3$ | 3.347 |
| ZINC74638808 | 346.4 | $C_{19}H_{26}N_2O_4$ | 1.859 |
| ZINC36391898 | 400.5 | $C_{25}H_{24}N_2O_3$ | 3.969 |
| ZINC30012975 | 424.5 | $C_{23}H_{24}N_2O_4S$ | 2.695 |
| ZINC12530852 | 493.7 | $C_{28}H_{35}N_3O_3S$ | 3.331 |
| ZINC05348237 | 379.5 | $C_{24}H_{29}N_1O_4$ | 1.359 |

*Final selection of MSM targets for virtual screening:* As aforementioned, MSM analysis determined that the most statistically prominent targets are A (17.3%), B (20.3%), and C (39.2%), with the statistical relevance of the remaining seven being at least an order of magnitude lower than that of target A (i.e. target H has 8.1% relevance). While target C is the most statistically significant, its docking region lacks the extraneous space extending from the allosteric pocket towards the A-loop, which are present in targets A and B. This morphological difference in the docking regions is reflected in target C’s docking volume of 1747 Å^3^, which is 20-30% smaller compared to the volume of the docking regions of targets A (2286 Å^3^) and B (2489 Å^3^). As previously discussed, this region initially seems extraneous, but the binding poses of Type III inhibitors PD318088^39^ and GDC-0973^40^ in MEK1/2 and p38α suggest that this region in MSM states of intermediate Src could be exploited for enhancing Src selectivity. Hence, using target C for virtual screening could restrict the chemical search space of lead Type III ligands selective to the allosteric pocket in Src extending towards the A-loop. Therefore, we considered candidates A and B as the two most statistically significant targets with docking regions best representing the majority of target conformations accessed during MD trajectories.

While target B is slightly more statistically significant than A, both have docking regions that are quite spacious and include the region extending from the allosteric pocket and towards the A-loop. However, that target A demonstrates better binding pose reproducibility for both ATP-Mg^2+^ and ANS, in addition to having a higher AUC value for the ATP-competitive region than target B (A: 0.722, B: 0.635) leads us to place slightly more confidence in the virtual screening results from target A. For these reasons, we used target A to screen the ZINC library of 13,195,609 ‘clean, drug-like’ ligands,^32^ followed by cross-docking the top-scoring ligands in target A against target B.

### Lead ligand enrichment of Type II and III allosteric inhibitors of Src

After using target A to screen the ZINC library of 13 million “clean, drug-like” ligands, we ranked all ligands in decreasing order by their Hammerhead score and selected the top 0.05% (6500 total) of highest-scoring ligands. We chose the top 0.05% because this cutoff is typically applied in virtual screening using ZINC libraries, as demonstrated in Kolb et al. for the β_2_-adrenergic receptor^41^. These 6500 ligands were clustered by their pairwise Tanimoto shape and chemical complementarity score (see Equation 1 in Text S2), using ROCS in the OpenEye Scientific Software^42^. (ROCS reports the Tanimoto score within a range from 0.0 to 2.0, representing a range of combinations of shape and chemical complementarity.) Figure S5 shows the histogram of all Tanimoto scores; the histogram peaks around 0.2, corresponding to 59 clusters. Each cluster is represented by a ‘centroid’ ligand with a Tanimoto score that collectively represents the shape, physicochemical properties, and binding poses of all ligands in the cluster. That the histogram peaks at 0.2 reflects the ample diversity in ligand predicted poses, which is due to the spaciousness of target A’s docking region. This clustering step allowed us to reduce the 6500 ligands to a smaller set of representative ligand binding poses that were subject to visual inspection, as follows.

The objective of lead enrichment is to identify Type II and Type III allosteric ligands that would displace the αC-helix outwards in a manner to prevent downstream Src activation. To get a general idea of where in Src do the 6500 ligands bind, we manually inspected their binding poses in target A. Consequently, we identified 500 ligands whose predicted binding pose in target A occupied the ATP-competitive and allosteric pocket (Type II) or strictly to the allosteric pocket (Type III). We then cross-docked the 500 ligands against target B; despite its caveats of lower binding pose reproducibility for ANS, target B is 3% more statistically significant than target A and exhibits a structurally similar allosteric pocket to that of target A. Since slight differences in the size and morphology of the docking regions in targets A and B could translate to differences in predicted poses of the 500 cross-docked ligands, we evaluated for each ligand’s binding pose consistency by superimposing the two target-ligand complexes and calculating the RMSD between the ligand’s poses in targets A and B. Ligands exhibiting an RMSD not exceeding 4 Å between the two targets were regarded as exhibiting excellent binding pose consistency. Accordingly, 250 ligands were identified as exhibiting excellent binding pose consistency in targets A and B. These ligands were then enriched based on whether they exhibited pharmacophoric features similar to ANS: hydrogen bonding to the Lys295 side chain amide and DFG backbone atoms and hydrophobic interactions between the ligand’s hydrophobic substructures and surrounding residues in the target (see Figure 1). LigandScout^43^ was used determine the pharmacophoric features for the 250 ligands bound to targets A and B.

### Performing FragFEATURE analysis on the MSM targets

The final step of lead enrichment is to prioritize lead ligands whose substructures are empirically supported by the FragFEATURE knowledgebase of protein pockets annotated with preferred fragment binding interactions. Using MSM targets A and B as our query targets, we determined the fragment-pocket binding preferences for the ATP-competitive and allosteric regions, represented by the residues within 5 Å of ATP-Mg^2+^ and ANS (see Table S5). For these residues, FragFEATURE calculates the microenvironments (single or multiple backbone and sidechain heavy atoms of a residue) and compares each microenvironment to a knowledgebase of microenvironments from proteins whose sequence identity to the query target is at most 50%.^25^ Each fragment prediction is characterized by a set of microenvironments and the set spread, or the maximum distance between two microenvironments of a microenvironment set (see Table S5). For each microenvironment set, FragFEATURE also calculates a Fishers’ p-value, or a probabilistic measure of fragment reliability; the value of the Fishers’ p-value is inversely related to the statistical significance of the predicted fragment. For MSM target A, FragFEATURE generated 371 fragments, of which 190 and 181 respectively correspond to the ATP-competitive and allosteric regions; for MSM target B, FragFEATURE predicted 501 fragments, of which 429 and 72 respectively correspond to the ATP-competitive and allosteric regions. These relatively high numbers of predicted fragments are reasonable given that the ATP-competitive and allosteric regions comprise a large composite binding pocket. Of all the fragment predictions for the two targets, those used for lead enrichment were fragments with a Fisher’s p-value of at least 10^-4^, set spread of at least 6.0, and having at least three microenvironment sets. Applying these criteria to the lead ligands excellent binding pose consistency, hydrogen bonding with Lys295’s side chain amide and DFG backbone amides, and ample hydrophobic interactions with surrounding Src residues, we identified sixteen allosteric lead ligands (seven Type II and nine Type III) with favorable fragment profiles, as shown in Figures 3 and 4.

**Figure 3.**
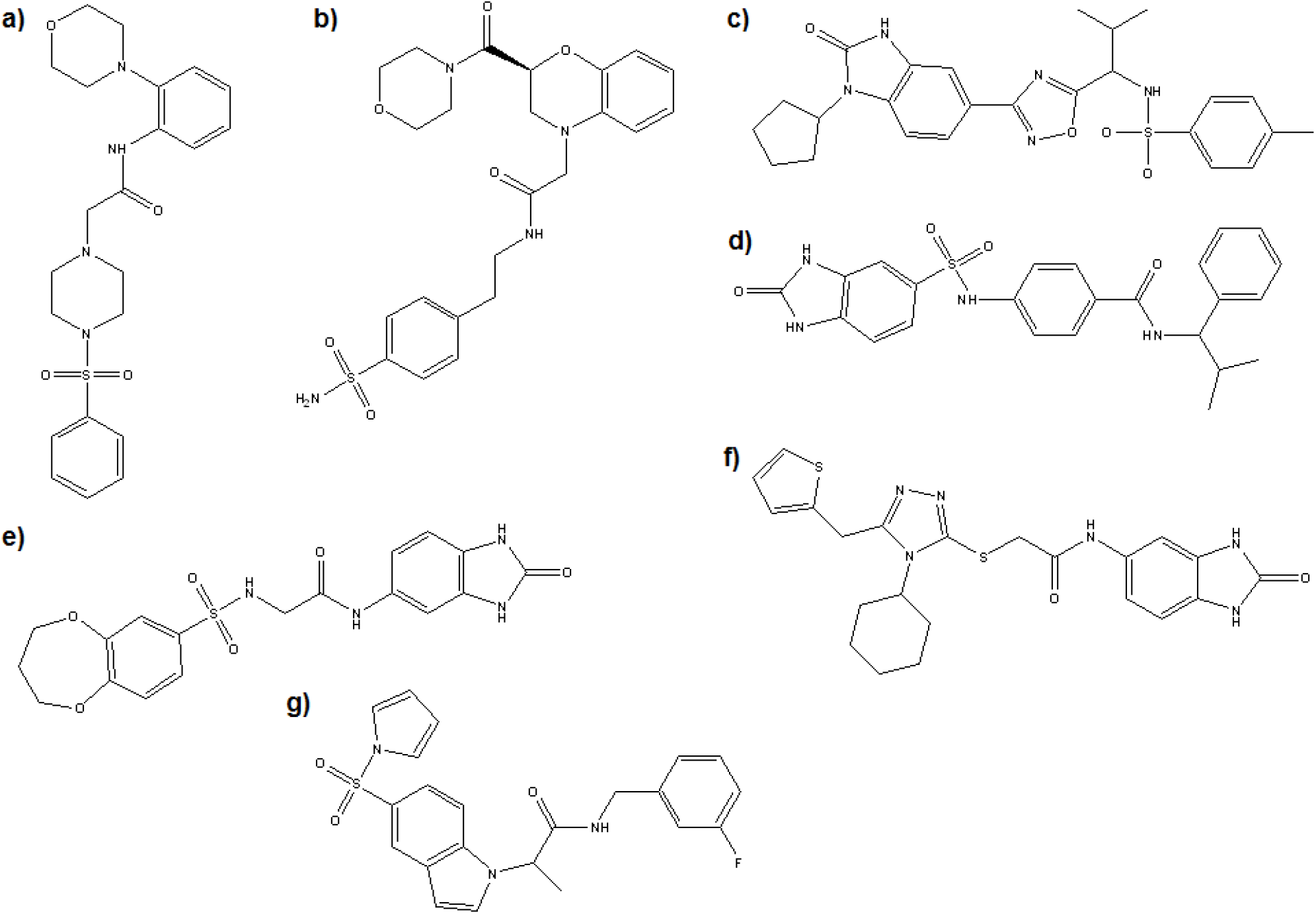
Chemical structures of Type II lead ligand candidates. Shows the structures for a) ZINC13037947, b) ZINC09672647, c) ZINC15729866, d) ZINC39795560, e) ZINC12939211, f) ZINC09125889, and g) ZINC09276644.

**Figure 4.**
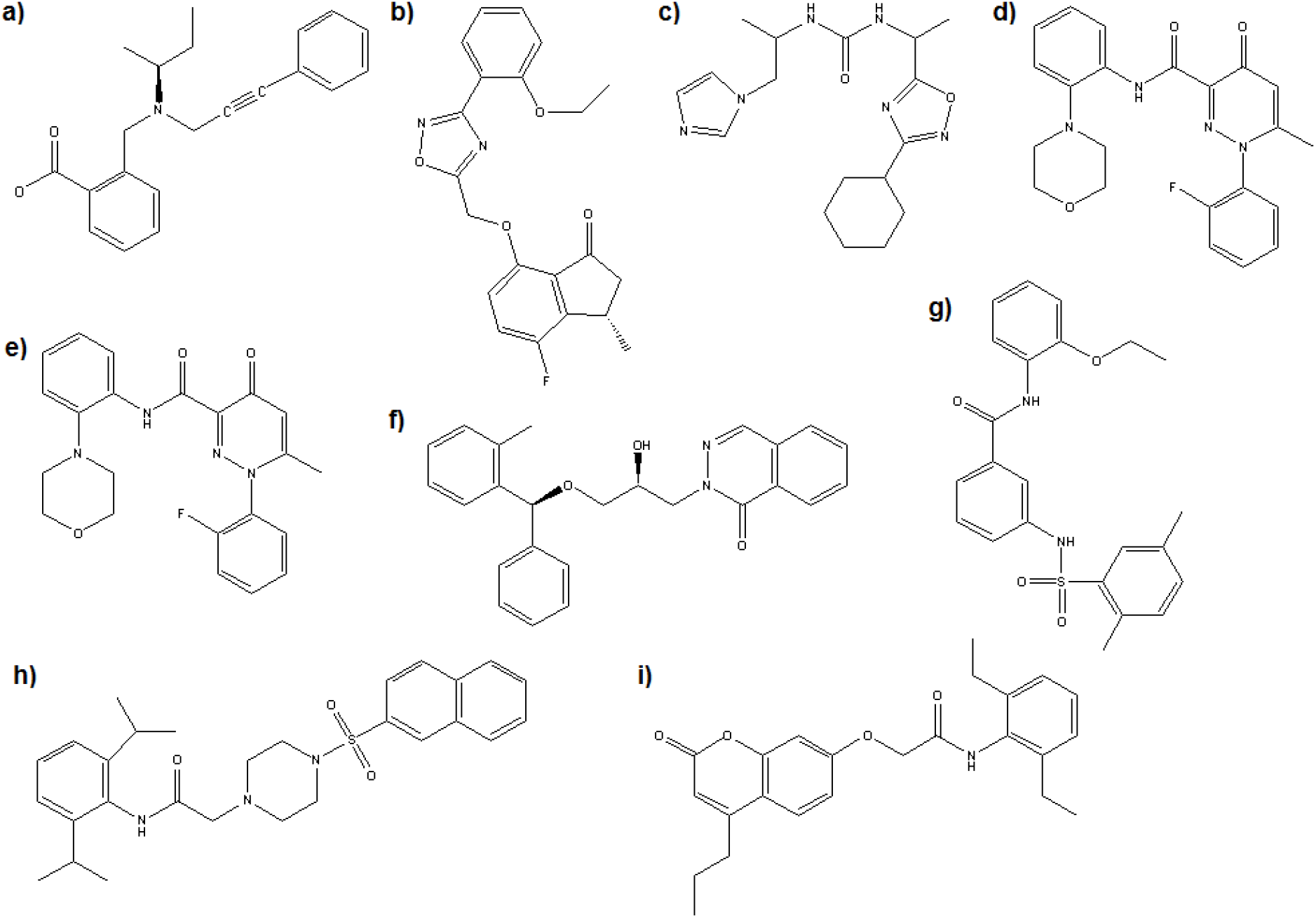
Chemical structures of Type III lead ligand candidates. Shows the structures for a) ZINC55183050, b) ZINC14369395, c) ZINC80660181, d) ZINC12543756, e) ZINC74638808, f) ZINC36391898, g) ZINC30012975, h) ZINC12530852, and i) ZINC05348237.

## RESULTS AND DISCUSSION

### Reserving FragFEATURE analysis as the final step in lead enrichment

Shukla et al.’s MD simulations of ANS-bound intermediate Src strongly suggest the existence of this allosteric pocket in presence of ANS. However, these results have not yet been experimentally validated, and there are also currently no known ligands that bind this allosteric pocket in a DFG-in conformation of Src. Therefore, in order to maximize the chemical search space of lead allosteric ligands of Src, we chose to screen a ZINC library of 13 million *known* ligands, rather than design a new ligand from fragments. FragFEATURE can be used as the first step of lead enrichment in fragment-based approaches, but this approach would be more suitable for protein pockets that have been well-studied such that selection of a starting fragment is decided using a wealth of historical information on pocket annotated with ligands. Even though the approach of enhancing kinase selectivity through targeting non-ATP-competitive regions has been gaining more traction in recent years, the allosteric region still has not been as intensely pursued as the ATP-competitive region. FragFEATURE even returned more fragments for the ATP-competitive region than for the allosteric region in MSM targets A and B. For these reasons, we reserved FragFEATURE as the very last lead enrichment step, after whole ligands have been prioritized.

### Characterization of pharmacophoric features and fragment profiles of Type II lead ligands

Our lead enrichment protocol identified seven Type II lead ligands (ZINC13037947, ZINC09672647, ZINC15729866, ZINC39795560, ZINC12939211, ZINC09125889, and ZINC09276644) exhibiting good binding pose consistency (RMSD < 4.0 Å) between MSM targets A and B (see Table 4). As shown in Figures 3 and S8-S14, the binding poses of these ligands in the two MSM targets could be stabilized by hydrogen bonding and hydrophobic interactions with various residues in the ATP-competitive and allosteric pocket of Src. Full descriptions of the Src residues involved in pharmacophoric interactions are provided in Tables S3a-b. These seven lead ligand’s poses in MSM targets A and B are further reinforced by FragFEATURE’s predicted fragments supporting substructures bound to the ATP-competitive and allosteric pockets of the MSM targets (see Table 4).

**Table 4.** Fragment profiles for Type II lead ligands. Descriptions of fragments and their corresponding microenvironments are provided in Table S5.

| ZINC ID | In MSM target A |  |  | In MSM target B |  |  |
| --- | --- | --- | --- | --- | --- | --- |
|  | No. distinct substructures supported by fragment predictions | Substructure: supporting fragment(s) | Total no. of fragment predictions | No. distinct substructures supported by fragment predictions | Substructure: supporting fragment(s) | Total no. of fragment predictions |
| ZINC13037947 | 2 | 4-phenylmorpholine: benzene, aniline, methylaniline<br>N,N-dimethyl methanesulfonamide : N-methyl methanesulfonamide | 4 | 3 | 4-phenylmorpholine: benzene, aniline, N-phenylformamide<br>N,N-dimethyl methanesulfonamide : N-methyl methanesulfonamide<br>trimethylamine: dimethylamine | 6 |
| ZINC09672647 | 2 | methylaniline: aniline, methylaniline<br>toluene: toluene | 3 | 3 | methylaniline: aniline<br>toluene: toluene<br>trimethylamine: dimethylamine | 4 |
| ZINC15729866 | 2 | N-methyl methanesulfonamide : N-methyl methanesulfonamide<br>toluene: toluene | 2 | 1 | N-methyl methanesulfonamide : N-methyl methanesulfonamide | 1 |
| ZINC39795560 | 1 | aniline: aniline | 1 | 2 | toluene: toluene<br>benzene: benzene | 2 |
| ZINC12939211 | 1 | benzene: benzene | 2 | 1 | mesylaniline: benzene, N-methyl methanesulfonamide | 2 |
| ZINC09276644 | 1 | fluorobenzene: fluorobenzene | 1 | 1 | fluorobenzene: fluorobenzene, N-methylbenzylamine | 2 |
| ZINC09125889 | 0 | n/a | 0 | 1 | dimethylamine: dimethylamine | 1 |

Evaluation of a ligand’s fragment profile was done based on the number of distinct substructures supported by FragFEATURE’s predicted fragments. Two substructures of a ligand were regarded as distinct in that their atoms in the ligand do not overlap. Accordingly, two predicted fragments that support the same substructure of a ligand also physically overlap. For example, the aniline substructure in ZINC39795560 is supported by aniline in MSM target A and two distinct substructures, toluene and benzene, in MSM target B (see Figure S11). ZINC09125889 is not supported by any fragment predictions in MSM target A and only one dimethylamine fragment in MSM target B (see Figure S12). The likelihood that a ligand has three distinct substructures supported by fragment predictions is lower than a ligand having only one or two distinct substructures supported by fragment predictions. Consequently, we regard ZINC13037947 as having the strongest fragment profile of all seven Type II lead ligands for its highest number of distinct substructures supported by FragFEATURE.

As shown in Figures 5 and S8, ZINC13037947 has two and three distinct substructures supported by fragment predictions in MSM targets A and B, respectively. The 4-phenylmorpholine substructure binding the allosteric pocket could be stabilized by hydrophobic interactions with Phe307, Ala311, Met314, Val323, Leu325, and Ile336 in both MSM targets A and B. Binding of 4-phenylmorpholine in target A is further reinforced by benzene, aniline, and methylaniline; binding of 4-phenylmorpholine in target B is supported by benzene, aniline, and N-phenylformamide. The binding of N,N-dimethylmethanesulfonamide could be stabilized by hydrogen bonding with Cys277 and Phe278 in MSM target A and only Phe278 in MSM target B and hydrophobic interactions with Val281 in both targets. The N,N-dimethylmethanesulfonamide substructure is also supported by N-methyl methanesulfonamide in both MSM targets A and B. Moreover, the trimethylamine substructure in MSM target B is also supported by dimethylamine fragments. In addition to having an excellent binding pose consistency of RMSD = 2.34 Å, ZINC13037947 is an excellent candidate of a Type II allosteric Src inhibitor.

**Figure 5.**
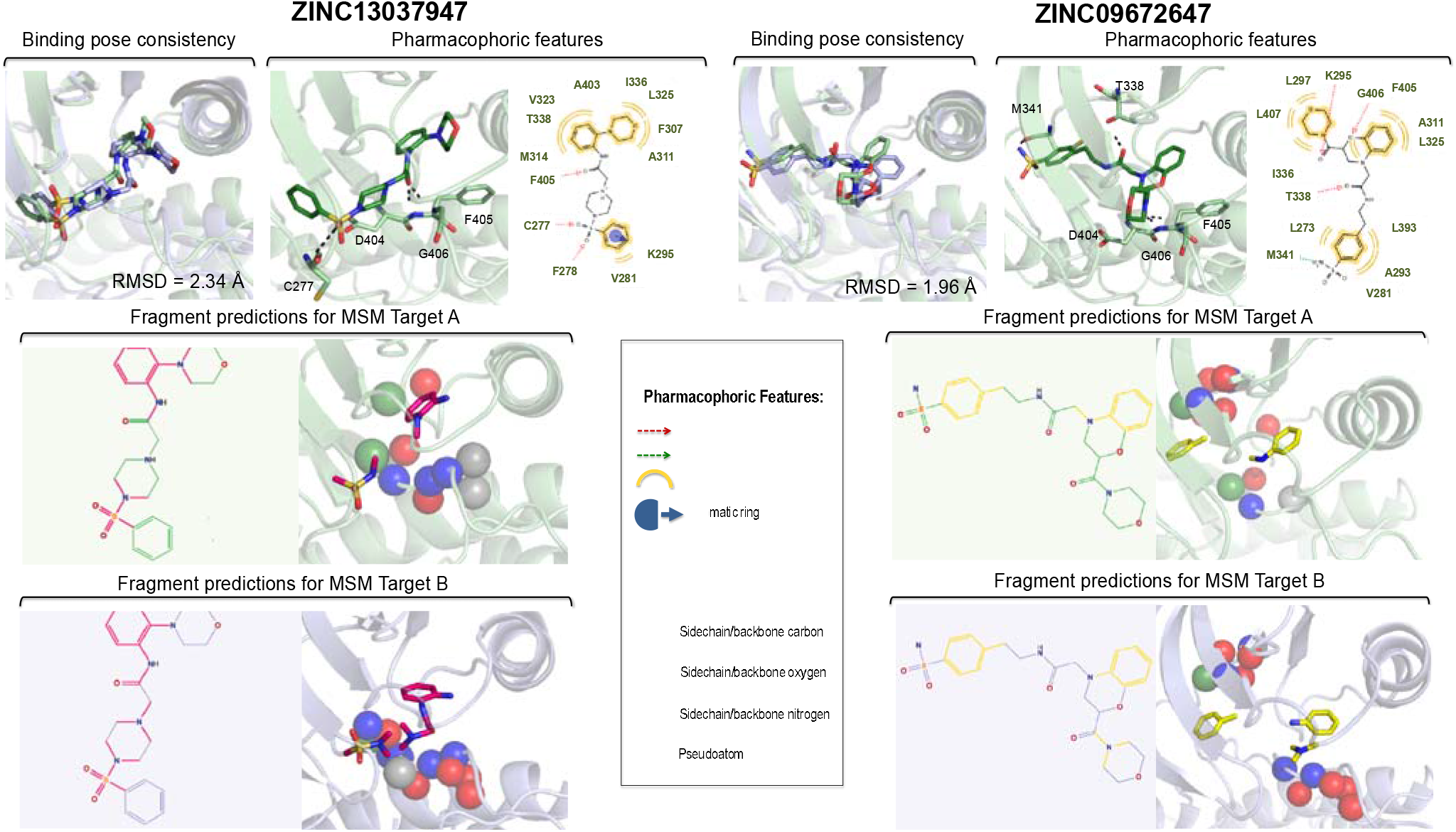
Type II allosteric inhibitors with strong fragment profiles. Information corresponding to MSM targets A and B are depicted in green and purple, respectively. Color schemes of hydrogen bonding, hydrophobic interactions, aromatic interactions, and FragFEATURE microenvironments are shown in the legend. The binding pose of ZINC13037947 in MSM target A can be stabilized by hydrogen bonding with Cys277, Phe278, and Phe405, hydrophobic interactions with Val281, Phe307, Ala311, Met314, Val323, Leu325, Ile336, Thr338, and Ala403, and an aromatic ring interaction with Lys295. This ligand’s binding pose in MSM target A is also reinforced by methylaniline, aniline, and benzene, all of which physically overlap, and N-methyl methanesulfonamide. The binding pose of ZINC13037947 in MSM target B is supported by benzene, aniline, N-phenylformamide, all of which three physically overlap, dimethylamine, and N-methyl methanesulfonamide. For ZINC09672647, its binding pose in MSM target A could be stabilized by hydrogen bonding with Lys295, Thr338, Gln339, Met341, and Gly406 and hydrophobic interactions with Leu273, Val281, Ala293, Lys297, Ala311, Leu393, and Phe405. This ligand’s binding pose in target A is supported by toluene and two physically overlapping fragments, methylaniline and aniline. The binding pose of ZINC09672647 in target B is supported by aniline, two overlapping dimethylamine fragments, and toluene.

Another Type II lead ligand with a strong fragment profile is ZINC09672647, which demonstrates an excellent binding pose consistency of RMSD = 1.96 Å. In MSM target A, the methylaniline substructure bound to the allosteric pocket is supported by aniline and methylaniline fragments; in MSM target B this substructure is only supported by aniline (see Figures 5 and S9). This methylaniline substructure could also be stabilized by hydrophobic interactions with Ala311, Leu325, and Phe405 in MSM target A and hydrophobic interactions with Met314, Leu325, and Phe405 in MSM target B. The toluene substructure bound to the ATP-competitive region is supported by toluene in both MSM targets and could also be stabilized by hydrophobic interactions with Leu273, Ala293, and Leu393 in both MSM targets. Two physically overlapping dimethylamine fragments supporting the trimethylamine substructure of ZINC09672647 in MSM target B (see Figure S9). Figures 5 and S9 also show that the binding pose of ZINC09672647 could be stabilized by hydrogen bonding between the sulfonamide’s oxygen and Met341, as well as hydrogen bonding with Lys295, Phe405, and Gly406. Hydrophobic interactions involving Leu273, Val281, Ala293, Lys297, Ala311, Leu393, and Phe405 in MSM target A and Leu273, Val281, Leu297, Phe307, Met314, Leu325, Leu393, and Leu407 in MSM target B also contribute to the ligand’s binding. In all, its strong pharmacophoric and fragment profiles deem a strong Type II Src lead candidate.

The fragment profiles of the other five Type II lead ligands are slightly weaker than those of ZINC39795560 and ZINC09672647 because they have only one or two substructures empirically supported in both or either MSM targets A and B, notably, ZINC15729866, ZINC39795560, ZINC12939211, and ZINC09276644 (see Table 4). Finally, ZINC09125889 has the weakest fragment profile because its binding pose in MSM target A lacks fragment validation and only one substructure is supported by FragFEATURE in MSM target B. Detailed descriptions of the pharmacophoric and fragment profiles of these five ligands are provided in the captions of Figures S10-S14.

### Characterization of pharmacophoric features and fragment profiles of Type III lead ligands

Our multi-stage lead enrichment protocol also identified nine Type III lead ligands (ZINC12530852, ZINC30012975, ZINC55183050, ZINC14369395, ZINC36391898, ZINC05348237, ZINC80660181, ZINC74638808, ZINC12543756) exhibiting good binding pose consistency (RMSD < 4.0 Å) between MSM targets A and B (see Table 5). As shown in Figures 3 and S15-S23, the binding poses of these ligands in the two MSM targets could be stabilized by hydrogen bonding and hydrophobic interactions with residues in the allosteric pocket of Src. Full descriptions of the Src residues involved in pharmacophoric interactions are provided in Tables S4a-b.

**Table 5.** Fragment profiles for Type III lead ligands. Descriptions of fragments and their corresponding microenvironments are provided in Table S6.

| ZINC ID | In MSM target A |  |  | In MSM target B |  |  |
| --- | --- | --- | --- | --- | --- | --- |
|  | No. distinct substructures supported by fragment predictions | Substructure and supporting fragment(s) | Total no. of fragment predictions | No. distinct substructures supported by fragment predictions | Substructure and supporting fragment(s) | Total no. of fragment predictions |
| ZINC12530852 | 2 | 2,6-dimethylaniline: benzene, N-methyl-o-toluidine, aniline<br>N-methyl methanesulfonamide: N-methyl methanesulfonamide | 4 | 3 | 2,6-dimethylaniline: benzene, toluene<br>N-methyl methanesulfonamide: N-methyl methanesulfonamide<br>trimethylamine: dimethylamine | 5 |
| ZINC30012975 | 1 | formanilide: benzene, N-phenylformamide | 2 | 2 | formanilide: benzene<br>mesylaniline: aniline, N-methylmethanesulfonamide | 3 |
| ZINC55183050 | 1 | benzene: benzene | 1 | 1 | benzene: benzene | 1 |
| ZINC14369395 | 1 | 4-fluoroanisole: benzene, fluorobenzene, anisole | 3 | 1 | 4-fluoroanisole: fluorobenzene | 1 |
| ZINC36391898 | 1 | benzene: benzene | 1 | 1 | benzene: benzene | 1 |
| ZINC05348237 | 1 | 2',6'-dimethylacetanilide: benzene, N-phenyl propanamide, aniline | 3 | 1 | 2',6'-dimethylacetanilide: benzene, toluene, N-methyl-o-toluidine | 3 |
| ZINC80660181 | 0 | n/a | 0 | 2 | N,N'-dimethyl methanediamine: dimethylamine (2)<br>dimethylamine: dimethylamine | 3 |
| ZINC74638808 | 0 | n/a | 0 | 1 | dimethylamine: dimethylamine | 1 |
| ZINC12543756 | 1 | N-(2-aminophenyl) formamide: aniline, N-phenyl formamide | 2 | 0 | n/a | 0 |

Based on the aforementioned criteria for the strength of a ligand’s fragment profile, ZINC12530852 has one of the strongest fragment profiles of all nine Type III ligand candidates, given that its number of distinct substructures (supported by fragment predictions) is the highest in both targets. Specifically, for ZINC12530852 bound to MSM target A, the 2,6-dimethylaniline substructure bound to the allosteric pocket is supported by benzene, N-methyl-o-toluidine, and aniline, all of which physically overlap (see Figures 6 and S15). Binding of 2,6-dimethylaniline could also be stabilized by hydrophobic interactions with Phe307, Ala311, Met314, Val323, Leu325, Ile336, Thr338, and Ala403. The sulfonamide substructure bound to the region extending towards the A-loop is supported by N-methyl methanesulfonamide and hydrogen bonding with Phe278. These two substructures are also supported in MSM target B: 2,6-dimethylaniline is supported by benzene and toluene; N-methyl methanesulfonamide also supports the sulfonamide substructure, in addition to hydrogen bonding with Lys295 and Phe278. Also, two overlapping dimethylamine fragments support the trimethylamine substructure in MSM target B. Overall, its ample pharmacophoric and fragment profiles make ZINC12530852 a strong Type III lead candidate.

**Figure 6.**
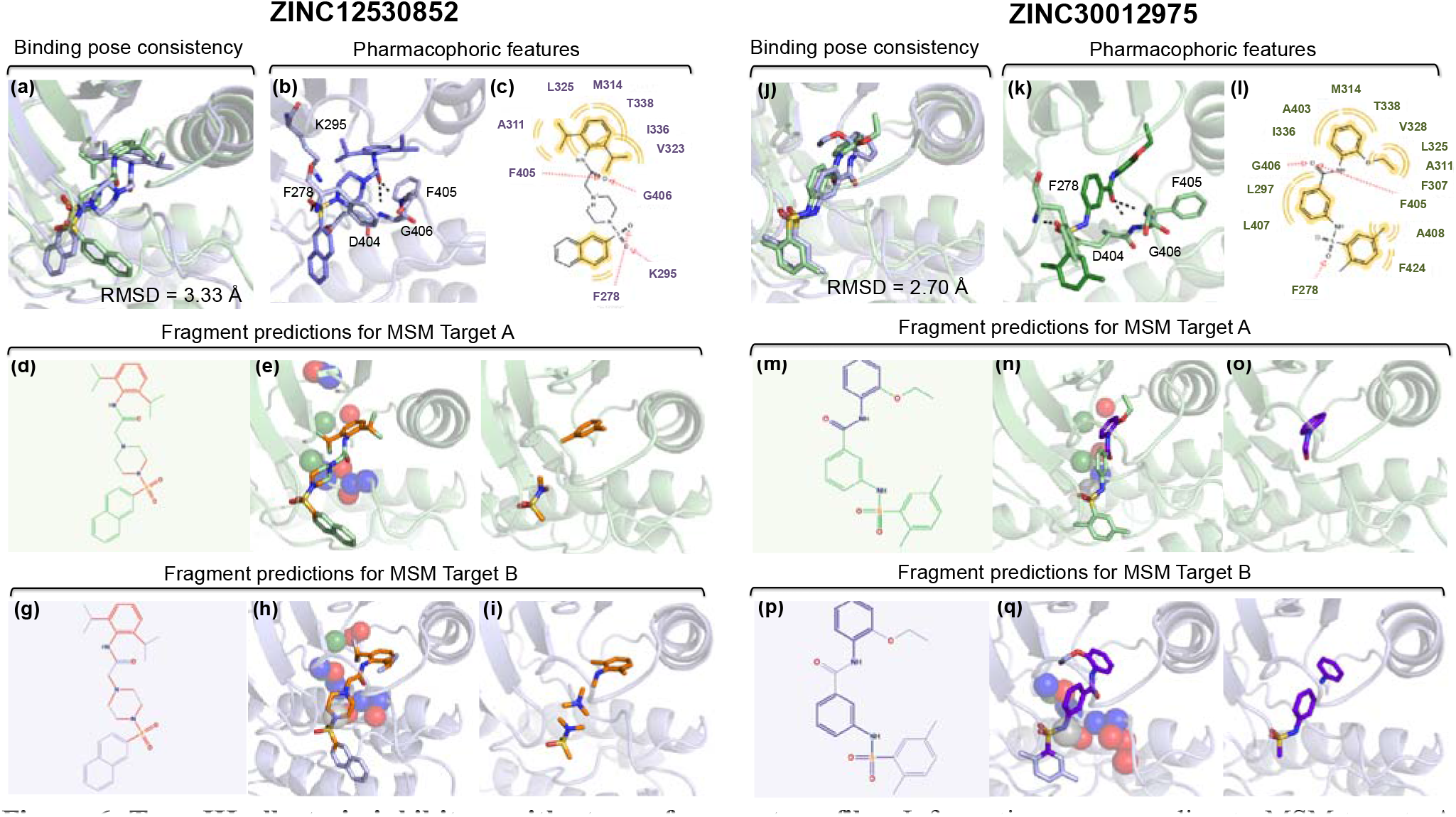
Type III allosteric inhibitors with strong fragment profiles. Information corresponding to MSM targets A and B are depicted in green and purple, respectively. The binding pose of ZINC12530852 in MSM target B can be stabilized by hydrogen bonding with Phe278, Lys295, Phe405, and Gly406 and hydrophobic interactions with Phe278, Ala311, Met314, Val323, Leu325, Ile336, and Thr338. This ligand’s binding pose in MSM target A is also reinforced by benzene, N-methyl-o-toluidine, and aniline, all of which physically overlap, and N-methyl methanesulfonamide. In target B, the binding pose of ZINC12530852 is supported by three physically overlapping fragment predictions, benzene, N-phenylformamide, and aniline, three overlapping dimethylamine fragments, and N-methyl methanesulfonamide. For ZINC30012975, its binding pose in MSM target A can be stabilized by hydrogen bonding with Phe278, Phe405, and Gly406 and hydrophobic interactions with Leu297, Phe307, Ala403, Ala311, Met314, Leu325, Val328, Ile336, Thr338, Leu407, Ala408, and Phe424. This ligand’s pose in target A is also supported by N-phenylformamide and benzene, which overlap, where as in target B, the binding pose is supported by three physically non-overlapping fragments, aniline, N-methyl methanesulfonamide, and benzene. Remaining pharmacophores and supporting fragment predictions for ZINC12530852 and ZINC30012975 are shown in Figures S15 and S16, respectively.

Another Type III lead ligand with a strong fragment profile is ZINC30012975, which has a formanilide substructure to the allosteric pocket and supported by benzene and N-phenylformamide; binding of formanilide could also be stabilized by hydrophobic interactions with Met314, Ile336, Thr338, and Ala403 and hydrogen bonding with Phe405 and Gly406 (see Figures 6 and S16). This formanilide substructure is supported by only benzene in MSM target B, and mesylaniline substructure is supported by aniline and N-methyl methanesulfonamide, along with hydrogen bonding with Phe278 and hydrophobic interactions with Lys295.

Of the seven other Type III lead candidates, four have only one substructure supported by FragFEATURE in both MSM targets, namely, ZINC55183050, ZINC14369395, ZINC36391898, and ZINC05348237, and thus have fragment profiles slightly weaker than those of ZINC12530852 and ZINC30012975. ZINC80660181, ZINC74638808, and ZINC12543756 have the weakest fragment profiles, as each ligand’s pose is supported by FragFEATURE in one but not both MSM targets. Detailed descriptions of the pharmacophoric and fragment profiles of these seven ligands are provided in the captions of Figures S17-S23.

## CONCLUSIONS

We have developed a target and lead enrichment methodology to identify sixteen Type II and III lead ligands towards confirming the existence of a potentially druggable allosteric pocket in intermediate Src conformations. Specifically, seven Type II and nine Type III lead ligands were identified to have excellent binding pose reproducibility and plentiful hydrogen bonding and hydrophobic interactions with Src residues. While these results suggest that these sixteen ligands should be pursued in experimental studies, FragFEATURE’s fragment predictions strengthened support for Type II ligands ZINC13037947 and ZINC09672647 and Type III ligands ZINC12530852 and ZINC30012975 as candidates with the strongest empirical evidence supporting their candidacy to inhibit Src activation. In all, we have demonstrated how MSM analysis can be used to identify statistically significant protein conformations for virtual screens, and show that the FragFEATURE knowledgebase can be used to enrich lead ligand candidates following virtual screening, through identifying leads with the best fragment profiles.

## ACKNOWLEDGEMENTS

This work was funded by the SIMBIOS NIH National Center on Biocomputing through the NIH Roadmap for Medical Research Grant U54 GM072970. M. L. is supported by the Ruth Kirschstein fellowship for postdoctoral researchers. D. S. is supported by the Biomedical Data Science Initiative postdoctoral scholar program of the Stanford School of Medicine. R. B. A is supported by LM05652, GM102365 and gifts from Microsoft, Oracle and Lightspeed Ventures. The computational resources for docking and chemotype clustering were performed on the Blue Waters supercomputer at the National Center for Supercomputing Applications at the University of Illinois at Urbana-Champaign and various clusters at Stanford University.

## REFERENCES

1 Kamb A, W. S., Lengauer C. Why is cancer drug discovery so difficult? Nat Rev Drug Discov 6, 115–120 (2007).

2 Venkatraman V, P.-N. V., Mavridis L, Ritchie DW. Comprehensive Comparison of Ligand-Based Virtual Screening Tools Against the DUD Data set Reveals Limitations of Current 3D Methods. J Chem Inf Model 50, 2079–2093 (2010).

3 F, S.-D. Target-based drug discovery: is something wrong? Drug Discov Today 10, 139–147 (2005).

4 Haspel N, M. M., Baker ML, Chiu W, Kavraki LE. Tracing conformational changes in proteins. BMC Struct Biol 10 (2010).

5 Boehr DD, N. R., Wright PE. The role of dynamic conformational ensembles in biomolecular recognition. Nat Chem Biol 5, 789–796 (2009).

6 al., S. D. e. Anton, a Special-Purpose Machine for Molecular Dynamics Simulation. Comm ACM 51, 91–97 (2008).

7 Kohlhoff KJ, S. D., Lawrenz M, Bowman GR, Konerding DE, Belov D, Altman RB, Pande VS. Cloud-based simulations on Google Exacycle reveal ligand modulation of GPCR activation pathways. Nature Chemistry 6, 15–21 (2013).

8 Hellerstein, J., Kohlkoff K, Konerding D. Science in the Cloud: Accelerating Discovery in the 21st Century. IEEE Internet Computing (2012).

9 Shirts M, P. V. Screen Savers of the World Unite! Science 290, 1903–1904 (2000).

10 Shan Y, A. A., Kim ET, Pan AC, Shaw DE. Transitions to catalytically inactive conformations in EGFR kinase. Proc Natl Acad Sci, doi:10.1073/pnas.1220843110 (2013).

11 Rucci N, S. M., Teti A. Inhibition of protein kinase c-Src as a therapeutic approach for cancer and bone metastases. Anticanc Agents Med Chem 8, 342–349 (2008).

12 Cowan-Jacob SW, F. G., Manley PW, Jahnke W, Fabbro D, Liebetanz J, Meyer T. The crystal structure of a c-Src complex in an active conformation suggests possible steps in c-Src activation. Structure 13, 861–871 (2005).

13 Roskoski Jr., R. Src protein-tyrosine kinase structure and regulation. Biochem Biophys Res Commun 324, 1155–1164 (2004).

14 Roskoski Jr., R. Src kinase regulation by phosphorylation and dephosphorylation. Biochem Biophys Res Commun 331, 1–14 (2005).

15 Aleshin, A., Finn, R. Src: a century of science brought to the clinic. Neoplasia 12, 599 (2010).

16 Yang S, R. B. Src Kinase Conformational Activation: Thermodynamics, Pathways, and Mechanisms PLoS Comp Biol 4, e1000047 (2008).

17 Betzi S, A. R., Martin M, Lubber DJ, Han H, Jakkaraj SR, George GI, Schönbrunn E. Discovery of a potential allosteric ligand binding site in CDK2. ACS Chemical Biology 6, 492–501 (2011).

18 Huang H, Z. R., Dickson BM, Skeel RD, Post CB. alphaC helix as a switch in the conformational transition of Src/CDK-like kinase domains. J Phys Chem B 116, 4465–4475 (2012).

19 Shukla D, M. Y., Roux B, Pande VS. Activation pathway of Src kinase reveals intermediate states as novel targets for drug design. Nat Comm (2014).

20 Adrián FJ, D. Q., Sim T, Velentza A, Sloan C, Liu Y, Zhang G, Hur W, Ding S, Manley P, Mestan J, Fabbro D, Gray NS. Allosteric inhibitors of Bcr-abl-dependent cell proliferation. Nat Chem Biol 2, 95–102 (2006).

21 Zhang J, Y. P., Gray NS. Targeting cancer with small molecule kinase inhibitors. 2009 9, 28–39 (2009).

22 J, B. Targeting tyrosine kinases in cancer: the second wave. Science 312, 1175–1178 (2006).

23 Lu S, H. W., Zhang J. Recent computational advances in the identification of allosteric sites in proteins. Drug Discov Today (2014).

24 Beauchamp, B. G., Lane TJ, Maibaum L, Haque IS, Pande VS. MSMBuilder2: modeling conformational dynamics on the picosecond to millisecond scale. J Chem Theory Comput 7, 3412–3419 (2011).

25 Tang GW, A. R. Knowledge-based fragment binding prediction. PLoS Comp Biol 10, e1003589 (2014).

26 Consortium, T. U. UniProt: a hub for protein information. Nucl Acids Res 43, D204–D212 (2014).

27 Shukla D, M. Y., Roux B, Pande VS. Activation pathway of Src kinase reveals intermediate states as novel targets for drug design. Nat Commun 5, doi:10.1038/ncomms4397 (2014).

28 Buch I, G. T., de Fabritiis G. Complete reconstruction of an enzyme-inhibitor binding process by molecular dynamics simulations. Proc Natl Acad Sci 108, 10184–10189 (2011).

29 Plattner N, N. F. Protein conformational plasticity and complex ligand-binding kinetics explored by atomistic simulations and Markov models. Nature Comm 6, doi:10.1038/ncomms8653 (2015).

30 Bowman GR, B. E., Hart KM, Maguire BC, Marqusee S. Discovery of multiple hidden allosteric sites by combining Markov state models and experiments. Proc Natl Acad Sci 112, 2734–2739 (2015).

31 Bowman GR, G. P. Equilibrium fluctuations of a single folded protein reveal a multitude of potential cryptic allosteric sites.Proc Natl Acad Sci 109, 11681–11686 (2012).

32 Irwin JJ, S. T., Mysinger MM, Bolstad ES, Coleman RG. ZINC: a free tool to discover chemistry for biology. J Chem Inf Model 52, 1757–1768 (2012).

33 Tang GW, A. R. Knowledge-based fragment binding prediction. PLOS Comp Biol 10 (2014).

34 AN, J. Surflex-Dock 2.1: robust performance from ligand energetic modeling, ring flexibility, and konwledge-based search. J Comput Aided Mol Des 26, 281–306 (2007).

35 Triballeau N, A. F., Brabet I, Pin J-P, Bertrand H-O. Virtual screening workfolow development guided by the “receiver operating characteristic” curve approach. Application to high-throughout docking on metabotropic glutamate receptor subtype 4. J Med Chem 48, 2534–2547 (2005).

36 Mysinger MM, C. M., Irwin JJ, Shoichet BK. Directory of useful decoys, enhanced (DUD-E): better ligands and decoys for better benchmarking. J Med Chem 55, 6582–6594 (2012).

37 Xu W, D. A., Lei M, Eck MJ, Harrison SC. Crystal structures of c-Src reveal features of its autoinhibitory mechanism. Mol Cell 3, 629–638 (1999).

38 Bissantz C, F. G., Rognan D. Protein-based virtual screening of chemical databases. 1. Evaluation of different docking/scoring combinations. J Med Chem 43, 4759–4767 (2000).

39 Ohren JF, C. H., Pavlovsky A, Whitehead C, Zhang E, Kuffa P, Yan C, McConnell P, Spessard C, Banotai C, Mueller WT, Delaney A, Omer C, Sebolt-Leopold J, Dudley DT, Leung IK, Flamme C, Warmus J, Kaufman M, Barrett S, Tecle H, Hasemann CA. Structures of human MAP kinase kinase 1 (MEK1) and MEK2 describe novel noncompetitive kinase inhibition. Nat Struct Mol Biol 11, 1192–1197 (2004).

40 Rice, K. D., Aay, N., Anand, N.K., Blazey, C.M., Bowles, O.J., Bussenius, J., Costanzo, S., Curtis, J.K., Defina, S.C., Dubenko, L., Engst, S., Joshi, A.A., Kennedy, A.R., Kim, A.I., Koltun, E.S., Lougheed, J.C., Manalo, J.C.L., Martini, J.F., Nuss, J.M., Peto, C.J., Tsang, T.H., Yu, P., Johnston, S. Novel Carboxamide-Based Allosteric Mek Inhibitors: Discovery and Optimization Efforts Toward Xl518 (Gdc-0973). ACS Med Chem Lett 3, 416 (2012).

41 Kolb P, R. D., Irwin JJ, Fung JJ, Kobilka BK, Shoichet BK. Structure-based discovery of β2-adrenergic receptor ligands. Proc Natl Acad Sci 106, 6843–6848 (2009).

42 ROCS, version 3.1.2 (Santa Fe, NM, 2011).

43 Wolber G, L. T. LigandScout: 3-D pharmacophores derived from protein-bound ligands and their use as virtual screening filters. J Chem Inf Model 45, 160–169 (2005).

